# Successful gene editing in tetraploid alfalfa using the open-source, AI-derived OpenCRISPR-1

**DOI:** 10.64898/2026.04.15.718787

**Authors:** Sameena Alam, Udaya Subedi, Kimberley Burton Hughes, Guanqun Chen, Letitia Da Ros, Andriy Bilichak, Stacy D. Singer

**Affiliations:** Agriculture and Agri-Food Canada, Lethbridge Research and Development Centre, Lethbridge, Alberta, T1J 4B1, Canada; Department of Agricultural, Food and Nutritional Science, University of Alberta, Edmonton, Alberta, T6G 2P5, Canada; Agriculture and Agri-Food Canada, Summerland Research and Development Centre, Summerland, British Columbia, V0H 1Z0, Canada; Agriculture and Agri-Food Canada, Morden Research and Development Centre, Morden, Manitoba, R6M 1Y5, Canada

**Keywords:** Alfalfa, Gene editing, OpenCRISPR-1, Plant biotechnology

## Abstract

While CRISPR/Cas-based gene editing technologies have the potential to greatly advance crop breeding endeavours, intellectual property-related challenges can hinder the ability to move such varieties to the market. Recently, an open-access Cas enzyme derived from large language models (OpenCRISPR-1) was developed and shown to function effectively in human cells. In this study, we demonstrate the successful use of this nuclease in a polyploid plant species (*Medicago sativa*), with mono- or bi-allelic editing observed in 30% of genotypes bearing OpenCRISPR-1. These findings indicate that OpenCRISPR-1 holds promise to expand the use of gene editing technology in the breeding of polyploid crops.

## Introduction

CRISPR/Cas platforms have become incredibly popular over the last decade to elicit mutations at specific genomic loci for a variety of purposes, providing unparalleled opportunities in medicine and crop breeding. Of CRISPR/Cas systems, those based on the Cas9 nuclease from *Streptococcus pyogenes* (SpCas9; Jinek et al. 2012) are currently the most commonly used in plants, allowing for target gene disruption, gene/allele replacement, base editing, prime editing, and epigenetic alterations, for instance (Singer et al. 2021). In many cases, CRISPR/Cas-induced mutations are indecipherable from those that occur spontaneously or through conventional breeding endeavours, and as such, the majority of nations have determined that crops derived from this type of editing will not face the same strict, lengthy and cost-prohibitive regulatory policies required for transgenic crops (Singer and Michaud 2025).

Unfortunately, however, the use of standard Cas-based technologies for crop breeding purposes is hindered substantially by a complex intellectual property (IP) landscape, which often involves multiple entities, enduring disputes, and inconsistencies among countries with respect to who holds the patents (Lin et al. 2025). Therefore, there is an urgent motivation to develop novel nucleases for gene editing purposes that are free of such restrictions. In line with this, generative AI large language models were recently harnessed to design a novel Cas nuclease (OpenCRISPR-1; Profluent Bio, Berkeley, CA) that bears over 400 nucleotide differences from SpCas9, and more than 180 from its closest natural homolog, which corresponds to an amino acid identity of no more than 86.3% (Ruffolo et al. 2025) and a lack of coverage by associated patents. Unlike other Cas enzymes, this nuclease has been made open-access for both research and commercial purposes, which has major implications for facilitating the use of CRISPR/Cas gene editing as a breeding tool. While OpenCRISPR-1 has been shown to exhibit comparable effectiveness to SpCas9 in human cells (Ruffolo et al. 2025), its application in plants is only now beginning to emerge (Das et al. 2026 [PREPRINT]). However, there are currently no reports regarding its efficacy in polyploid or dicotyledonous plant species, which make up a substantial proportion of crops. Therefore, we sought to assess its ability to elicit mutations at a test locus (*PLANTACYANIN4*; *MsPLC4*) in the important tetraploid forage legume, alfalfa (*Medicago sativa* L.), as a means of validating this potentially transformative editing tool for inclusion in our crop breeding toolbox.

## Materials and methods

### Development of gene editing vectors

To assess the efficacy of the OpenCRISPR-1 system in alfalfa, the *MsPLC4* gene was targeted using both SpCas9-(positive control) and OpenCRISPR-1-based vectors. The OpenCRISPR-1 coding sequence (https://github.com/Profluent-AI/OpenCRISPR) was codon-optimized for *Triticum aestivum* (primary expression host) and *Medicago sativa* (secondary expression host), and translationally fused at both 5’ and 3’ ends to nuclear localization signals (NLS). The resulting translational fusion was synthesized by a service provider (GenScript Biotech Corp., Piscataway, NJ; File A1). In the case of the SpCas9 control, the pKSE401 vector (Xing et al. 2014; Addgene plasmid #62202), which includes a translational fusion comprising 3x FLAG - NLS - *Zea mays* codon-optimized SpCas9 coding sequence - NLS, was used. To enable comparison between the two nucleases, the SpCas9 translational fusion in pKSE401 was replaced with the OpenCRISPR-1 translational fusion using restriction-based cloning to generate the pKSE401-OpenCR vector.

A Basic Local Alignment Search Tool (BLAST) alignment of the *PLC4* coding sequence from *Medicago truncatula* (Medtr8g08910), which is a close diploid relative of autotetraploid alfalfa with a very well-annotated genome sequence, was conducted against the reference genome of a genotype of XinJiangDaYe alfalfa (Chen et al. 2020). Three guide RNAs (gRNAs; 20 nt) immediately upstream of 5’ – NGG – 3’ protospacer adjacent motifs (PAMs) were selected using CRISPR-P 2.0 software (http://crispr.hzau.edu.cn/cgi-bin/CRISPR2/CRISPR; Liu et al. 2017) in exons 1 and 2 of the *MsPLC4* genomic sequence that were identical across all four alleles (Fig. 1A). Golden Gate cloning was utilized as described previously (Xing et al. 2014) to assemble the three gRNAs with an SpCas9 single gRNA scaffold. This was then inserted into the SpCas9- (pKSE401) and OpenCRISPR-1-based (pKSE401-OpenCR) background vectors (see Table A1 for primer sequences). The identities of the resulting vectors (termed PLC4-Cas9 and PLC4-OpenCR; Fig. 1B) were confirmed via sequencing.

**Figure 1.**
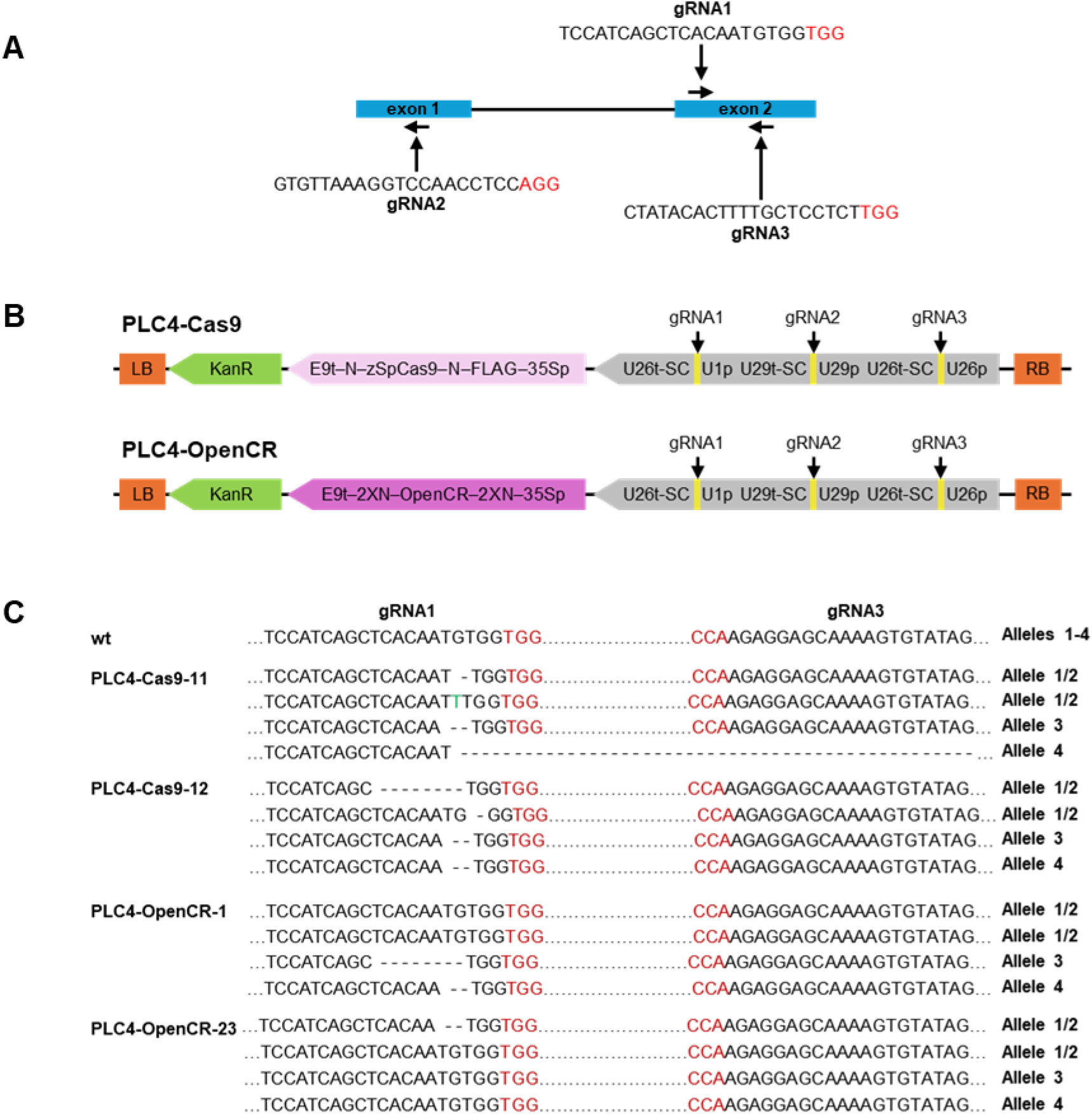
OpenCRISPR-1-mediated editing at the *MsPLC4* locus in alfalfa. **(A)** Genetic structure (not to scale) of *MsPLC4* and location of gRNAs within conserved exonic regions. Exons are denoted by blue boxes and the intron is indicated by a black line. **(B)** Diagrammatic representations (not to scale) of PLC4-Cas9 and PLC4-OpenCR vectors. 35Sp, partially duplicated CaMV 35S promoter; FLAG, 3x FLAG polypeptide tag; KanR, kanamycin resistance cassette including 35Sp, neomycin phosphotransferase II coding sequence, and nopaline synthase transcriptional terminator; LB, left border; N, nuclear localization signal; OpenCR, *OpenCRISPR-1* codon-optimized for *Triticum aestivum* (primary expression host) and *Medicago sativa* (secondary expression host); RB, right border; E9t, pea RuBisCO small subunit E9 terminator; SC, sgRNA scaffold; U26p, Arabidopsis U6-26 promoter; U26t, U6-26 terminator; U29p, Arabidopsis U6-29 promotor; U29t, U6-29 terminator; U1p, Arabidopsis U6-1 promotor; zSpCas9, *Zea mays* codon-optimized *SpCas9*. **(C)** Examples of PLC4-Cas9 and PLC4-OpenCR genotypes with precise mutations observed across alleles. Alleles 1 and 2 were identical in the wild-type (wt) genotype and made up approximately 50% of total reads. Dashes indicate deletions, nucleotides in red indicate PAM sequences, and nucleotides in green denote insertions.

### Alfalfa transformation

Both editing vectors were introduced into *Agrobacterium tumefaciens* strain LBA4404 via electroporation and were subsequently transformed into the alfalfa N4.2.2 genotype as described previously (Subedi et al. 2023). DNA was subsequently extracted from the leaves of 10 independent PLC4-Cas9 and PLC4-OpenCR transformants, respectively, as well as the untransformed alfalfa genotype, using the Biosprint 96 DNA Plant Kit (Qiagen, Germantown, MD). PCR was conducted using primers Cas9F1 and Cas9R1, or OpenCRISPRF1 and OpenCRISPRR1 (Table A1), respectively, along with Platinum SuperFi II Green PCR Master Mix according to the manufacturer’s instructions (Thermo Fisher Scientific, Nepean, ON) to verify the transgenic nature of each genotype.

### Amplicon sequencing

To identify potential mutations at *MsPLC4* target loci, 10 independent PLC4-Cas9 and PLC4-OpenCR genotypes, respectively, as well as the untransformed control, were assessed using a dual-indexed amplicon sequencing approach. A genomic region (568 bp) encompassing all three gRNA target sites was initially amplified (25 cycles) from approximately 40 ng genomic DNA from all genotypes using 0.2 μM of primers PLC4MiseqF1 and PLC4MiseqR1 (Table A1), along with Platinum SuperFi II Green PCR Master Mix in a total reaction volume of 25 μL according to the manufacturer’s recommendations (Thermo Fisher Scientific). A second round of PCR was conducted in the same manner using 5 μL of a 1/20 dilution of the original PCR amplicon as template, along with 0.2 μM of indexed forward (500 series) and reverse (700 series) primers (Table A1). The resulting PCR amplicons were purified using AMPure XP magnetic beads (Beckman Coulter Canada LP, Mississauga, ON) and paired-end sequencing (300 bp) was carried out using an Illumina MiSeq i100 platform (Illumina Inc., San Diego, CA) through a service provider (Integrated Microbiome Resource, Dalhousie University, Halifax, NS). The resulting reads were analyzed using the CRISPR analysis tool in Geneious Prime 2026.0.2 (GraphPad Software, LLC, Boston, MA), whereby reads were trimmed for quality using the BBDuk plugin (reads smaller than 50 nt were discarded), further trimmed to 250 nt, and those exhibiting a percent read value of less than 6.5% were excluded.

## Results and discussion

Four *MsPLC4* alleles were identified in the genome of tetraploid alfalfa (Chen et al. 2020; chr4.1 11588717 - 11589362, chr4.2 12366805 - 12367449, chr4.3 11250577 - 11251203, and chr4.4 12031295 - 12031921), which exhibited 98.4% nucleotide identity among the four alleles, as well as 97.3-97.8% nucleotide identity and 100% amino acid identity with the *M. truncatula PLC4* coding sequence and predicted protein sequence, respectively. Three gRNAs were designed to target all *MsPLC4* alleles simultaneously (Fig. 1A) and were inserted into the positive control SpCas9-based pKSE401 background vector (Xing et al., 2014) to generate PLC4-Cas9 (Fig. 1B). The gRNAs were also inserted into the experimental vector, pKSE401-OpenCR, which was identical to pKSE401 with the exception of the SpCas9 translational fusion, which had been replaced with a translational fusion including the *OpenCRISPR-1* coding sequence (PLC4-OpenCR; Fig. 1B).

Both vectors were introduced into alfalfa, and ten independent transgenic genotypes bearing each vector were selected for analysis. As expected, amplicon sequencing of a region of *MsPLC4* spanning all three gRNA sites indicated the presence of four *MsPLC4* alleles (two of which were identical, with the remaining two being distinct) in the untransformed wild-type genotype, with no evidence of mutations at any gRNA site. Also in line with expectations was the fact that mutations were observed within gRNA target sites in 80% of transgenic genotypes bearing the PLC4-Cas9 cassette, with 75% of these exhibiting mutations only at the gRNA1 site and 25% bearing mutations at both gRNA1 and gRNA3 sites (Fig. 1C; Table 1). Interestingly, we also observed on-target editing (at the gRNA1 site only) in 30% of independent transgenic genotypes bearing the PLC4-OpenCR cassette. No edits were detected at the gRNA2 site in either case. This difference in gRNA efficiency is commonly observed in plants (e.g., Wolabu et al. 2024). While the precise reasons for such divergences remain elusive, it is possible that epigenetic characteristics at the gRNA target site (e.g., Weiss et al. 2022), gRNA properties (e.g., Liang et al. 2016), or efficacy of the particular promoter used to drive gRNA expression (e.g., Zhu et al. 2023) could be contributing factors.

**Table 1.** Mutation type in PLC4-Cas9 and PLC4-OpenCR genotypes.

| Genotype | Alleles 1/2* | Allele 3 | Allele 4 | Mutant type |
| --- | --- | --- | --- | --- |
| wt | wt<br>wt | wt | wt | N/A |
| **PLC4-Cas9** |  |  |  |  |
| 3 | wt<br>1 bp del | 12 bp del | 2 bp del | Tri-allelic |
| 4 | wt<br>wt | wt | 1 bp del | Mono-allelic |
| 6 | 10 bp del<br>1 bp ins | no amplicon | 1 bp sub | Minimum tri-allelic |
| 7 | 9 bp del<br>5 bp del | 1 bp del at gRNA1 and 1 bp del at gRNA3 | no amplicon | Minimum tri-allelic |
| 8 | 1 bp del<br>no amplicon | wt | 1 bp insertion | Minimum bi-allelic |
| 11 | 1 bp del<br>1 bp ins | 2 bp del | 65 bp del between gRNA1 and gRNA3 | Tetra-allelic |
| 12 | 8 bp del<br>1 bp del | 2 bp del | 2 bp del | Tetra-allelic |
| 15 | 1 bp ins<br>no amplicon | 3 bp del | wt | Minimum bi-allelic |
| **PLC4-OpenCR** |  |  |  |  |
| 1 | wt<br>wt | 8 bp del | 2 bp del | Bi-allelic |
| 23 | wt<br>2 bp del | wt | wt | Mono-allelic |
| 24 | wt<br>1 bp del | wt | wt | Mono-allelic |

\*Alleles 1 and 2 were identical in the N4.4.2 wild-type genotype and made up approximately 50% of total reads. Del, deletion; ins, insertion; sub, substitution; wt, wild-type.

Of the four SpCas9-edited genotypes where four *MsPLC4* alleles were identified, one bore a mono-allelic mutation in *MsPLC4*, one possessed a tri-allelic mutation, and two bore tetra-allelic mutations with no wild-type sequence (Table 1), which aligns with previous studies in which SpCas9 has been utilized in alfalfa to obtain tetra-allelic mutations in the first generation (e.g., Wolabu et al. 2024). Furthermore, four other genotypes also possessed mutations in two to three *MsPLC4* alleles; however, only three alleles were detected in each case, suggesting the possibility of larger-scale alterations in one allele that may have prevented amplification (Table 1). Conversely, in the case of the three OpenCRISPR-1-edited genotypes, mono-allelic mutations were observed in two of the genotypes, while the third genotype bore bi-allelic mutations (Table 1). Although this apparent reduction in editing efficiency differed from the comparable activity of OpenCRISPR-1 and SpCas9 observed previously in human cells (Ruffolo et al. 2025), a slight decrease in mutation frequencies was also noted in regenerated rice plants in a preliminary report (Das et al. 2026 [PREPRINT]). While the more substantial reduction in efficiency seen in alfalfa could, at least in part, stem from codon optimization issues in our *OpenCRISPR-1* sequence, it will be imperative to carry out larger scale studies to gain further insight into the precise differences in efficacy between nucleases.

As is typical for SpCas9, all mutations in PLC4-Cas9 genotypes occurred 0-5 bp upstream of the PAM, with the majority consisting of 1-12 bp deletions (present in all eight genotypes), and one genotype also bearing a larger 65 bp deletion between gRNA1 and gRNA3 sites (Fig. 1C; Table 1). In addition, four genotypes also exhibited a 1 bp insertion, and one genotype bore a G to A substitution at the gRNA1 site (Fig. 1C; Table 1). Similarly, mutations observed in PLC4-OpenCR genotypes consisted of 1-8 bp deletions 3 bp upstream of the PAM sequence (Fig. 1C; Table 1), which corresponds with previous findings using OpenCRISPR-1 in human cells (Ruffolo et al. 2025).

## Conclusion

While a very small number of CRISPR-derived crop varieties, including γ-aminobutyric acid (GABA)-enriched ‘Sicilian Rouge’ tomatoes (Sanatech Seed) and non-browning lettuce (GreenVenus) have been released commercially in certain countries in recent years, the expense and convolution of licensing is limiting to many entities (Glenna 2023). In this study, we determined that the open-access OpenCRISPR-1 platform can successfully elicit editing in a tetraploid dicotyledonous plant species. While the efficiency of this system may be somewhat lower than SpCas9 in alfalfa, further research will be required to confirm this. In any case, the addition of this IP-free gene editing tool to our breeding toolbox for polyploid crops could open the door for a far wider range of both public and private institutions. This would facilitate the provision of enhancements in a more comprehensive suite of traits than what has been made available thus far, at least some of which could be geared towards public good.

## CRediT authorship contribution statement

**SDS:** Conceptualization, Funding acquisition, Methodology, Supervision, Writing – original draft, Writing – review and editing. **AB:** Conceptualization, Funding acquisition, Methodology, Writing – review and editing. **LDR:** Conceptualization, Writing - review and editing. **GC:** Supervision, Writing – review and editing. **KBH:** Formal analysis, Methodology, Writing – review and editing. **US:** Investigation, Formal analysis, Methodology, Writing – review and editing. **SA:** Investigation, Formal analysis, Methodology, Writing – review and editing.

## Data availability

The raw reads generated in this project are available at the National Center for Biotechnology Information (NCBI) Sequence Read Archive (BioProject accession number PRJNA1426072).

## Appendices

**Table A1**. Primer sequences

**File A1**. *OpenCRISPR-1* fusion sequence used in the generation of the alfalfa pKSE401-OpenCR vector

