## Supplementary material for "Successful gene editing in tetraploid alfalfa using the open-source, AI-derived OpenCRISPR-1": Table A1 and File A1

**Table A1**. Primer sequences

| **Primer name** | **Sequence (5’-3’)** |
| --- | --- |
| **Vector construction** | |
| PLC4gRNA1-BsF | ATATATGGTCTCGATTG**TCCATCAGCTCACAATGTGG**GTT |
| PLC4gRNA1-F0 | TG**TCCATCAGCTCACAATGTGG**GTTTTAGAGCTAGAAATAGC |
| PLC4gRNA2-BsF2 | ATATTATTGGTCTCAAGATTG**GTGTTAAAGGTCCAACCTCC**GTT |
| PLC4gRNA2-F0 | TG**GTGTTAAAGGTCCAACCTCC**GTTTTAGAGCTAGAAATAGC |
| PLC4gRNA3-R0 | AAC**AGAGGAGCAAAAGTGTATAG**CAATCACTACTTCGTCTCTAACCAT |
| PLC4gRNA3-BsR | ATTATTGGTCTCGAAAC**AGAGGAGCAAAAGTGTATAG**C |
| DT0-BsR2 | ATATTATTGGTCTCAATCTCTTAGTCGACTCTACCAAT |
| **Verification of transgenic genotypes** | |
| Cas9F1 | GCTGGAGGAGTCATTCCTCG |
| Cas9R1 | CTGAGAATATCAGACAGGAGG |
| OPENCRISPRF1 | GGTTTAAGGAGCATGAGGAAGA |
| OPENCRISPRR1 | TTCGAAGTTCCATGGTCTGATAG |
| **Amplicon sequencing** | |
| PLC4MiseqF1 | *TCGTCGGCAGCGTCAGATGTGTATAAGAGACAG*GCATTGGTTCTACTAGTCTGCT |
| PLC4MiseqR1 | *GTCTCGTGGGCTCGGAGATGTGTATAAGAGACAG*TGAAGTAGTTTTGTCCCCTAGC |
| Miseq-S508 index | *AATGATACGGCGACCACCGAGATCTACAC*CTAAGCCTTCGTCGGCAGCGT*C |
| MiSeq-S510 index | *AATGATACGGCGACCACCGAGATCTACACC*GTCTAATTCGTCGGCAGCGT*C |
| Miseq-S511 index | *AATGATACGGCGACCACCGAGATCTACAC*TCTCTCCGTCGTCGGCAGCGT*C |
| Miseq-S515 index | *AATGATACGGCGACCACCGAGATCTACAC*TTCTAGCTTCGTCGGCAGCGT*C |
| Miseq-S516 index | *AATGATACGGCGACCACCGAGATCTACAC*CCTAGAGTTCGTCGGCAGCGT*C |
| Miseq-S517 index | *AATGATACGGCGACCACCGAGATCTACAC*GCGTAAGATCGTCGGCAGCGT*C |
| Miseq-S518 index | *AATGATACGGCGACCACCGAGATCTACAC*CTATTAAGTCGTCGGCAGCGT*C |
| Miseq-S520 index | *AATGATACGGCGACCACCGAGATCTACAC*AAGGCTATTCGTCGGCAGCGT*C |
| Miseq-N724 index | *CAAGCAGAAGACGGCATACGAGAT*CGCTCAGTGTCTCGTGGGCTCG*G |
| Miseq-N726 index | *CAAGCAGAAGACGGCATACGAGAT*GTCTTAGGGTCTCGTGGGCTCG*G |
| Miseq-N727 index | *CAAGCAGAAGACGGCATACGAGAT*ACTGATCGGTCTCGTGGGCTCG*G |
| Miseq-N728 index | *CAAGCAGAAGACGGCATACGAGAT*TAGCTGCAGTCTCGTGGGCTCG*G |

gRNA sequences are indicated in bold, adapter sequences are indicated in italics, indices are underlined, and asterisks denote phosphorothioate bonds.

**File A1.** *OpenCRISPR-1* fusion sequence used in the generation of the alfalfa pKSE401-OpenCR vector

ATGCCTGCCGCCAAGCGTGTTAAGCTGGATGGAGGGAAAAGAACCGCCGATGGGTCAGAGTTTGAGTCACCTAAGAAAAAGAGGAAAGTTATGGTGAAGAAGCCATACTCCATTGGTTTGGACATCGGAACCAATAGCGTTGGCTGGGCAGTGATTACTGATGACTACAAGGTTCCTGCTAAGAAGATGAAGGTGCTTGGTAACACCGATAGGTCACATATTAAGAAGAATTTGATCGGAGCACTCCTTTTCGACGCCGGCAACACTGCAGAGGATAGGAGACTCAAGAGGACAGCTAGGAGAAGGTACACCAGAAGGAGAAACAGAATCTTGTACCTGCAAGAAATCTTCGCAGAGGAAATGAACAAGATCGACGAGTCCTTCTTTCACAGGCTCGATGACAGCTTTCTTGTTCCAGAAGATAAGAGAGGAAGCAAGTACCCTATCTTTGCTACACTTCAAGAGGAAAAGGAGTACCATAAGCAGTTCCCAACCATTTACCACCTCAGGAAGCAACTTGCCGAATCTAACGAGAAGGCAGATCTCAGACTTGTTTACTTGGCCCTGGCACATATGATCAAGTACAGGGGTCACTTCCTTATTGATGACCCTAAGTTTAAGGTGCAAAACAATGACATCCAGGGATTGTTCGAAAAGTTTGTTGAGGAATACGATAATGTGCAGGAGACTTCTCTCTCAAAGATTAAGCTGAACGTGACCGAAATTCTCACTGCCAAGATCCCAAAGTCCGAAAAGCAAGAGCAGTTGCTGAAGAATTACCCTAGCGAGAAGAAGAACACTTTGTTCGGCAATCTGATCGGTCTTGCCCTTGGATTGACCCCAAACTTCAAGACTAACTTCTCCTTGGAAAACGACGCAAAGCTGCAAATTTCTTCAGAGTCTTACGAGGAAGATCTCGGTTCACTCCTTGCTCTTATCGGAGAAAACTTCATCGAGTTGTTTTCTGCCGTGAAGAACCTGTCAGACGGCATTTTGCTGGCCGGTATCGTTTCCGATGAAAGCCCACATGCACCTTTGAGCACTAAGATGGTTATCAGGTTTAAGGAGCATGAGGAAGACCTGGCTGCCCTCAAGCACTTCATTAAGGCAAATCTCCCTGAAAAGTACGATGAGGTTTTCTCTGATGACTCAAAGAACGGATACGCTGGCTACGTTGGTGTGGATTCTAAAGTGAGGAAGAGAAATGGCAAGCTGGCCACAGAGGAAGAGTTCTACAAGTACCTCAAGGACATCCTTAACAATGTTAAGGGTGCCGATTACTTCCTTGAAAAGATTAAGAGAGAGGACCTCCTTAGGAAGCAAAGAACATTTGATAACGGAACCATCCCATACCAGGTGCATCTCGAAGAGATGAAGGCAATTCTTCAAAATCAGGGCGAGTACTACCCTTTCCTCAAGGAAAACAAGGAGAAGATTCAACAGATCCTTACATTTAGGATTCCATACTACGTTGGACCTCTCGCTAGGGGCAACAGAGACTTTGCCTGGCTTACCAGGAATTCAGATCAAGCTATCAGACCATGGAACTTCGAAGAGGTTGTGGACAAGGCTTCCAGCGCCGAGGATTTCATCAACAAGATGACTAACTACGATCTGTACCTCCCAGAAGAGAAGGTGCTGCCTAAGCACTCATTGCTGTACGAAACTTTCGCAGTTTACAATGAGCTTACAAAGGTGAAGTTCATCGCTGAGGGATTGAGAGACTACCAATTTCTGGATTCCGGCCAAAAGAAGCAGATTGTTAACCAGCTTTTTAAGGAAAAGAGGAAAGTGACTGAGAAGGATATTATCCATTACTTGCACAATGTTGACGGATACGATGGCATTGAACTGAAGGGTATCGAGAAGCAATTCAACGCATCCCTCAGCACATACCATGACCTCCTTAAGATCATCAAGGATAAGGAATTCATGGATGACCCAAAGAACGAAGAGATTTTGGAGAATATCGTTCACACTCTGACAATCTTCGAAGATAGGGAGATGATTAAGCAAAGACTCGCTCAGTACGACTCACTTTTTGATGAGAAAGTGATTAAGGCCCTTACTAGGAGACACTACACAGGCTGGGGAAAGTTGTCCGCTAAGCTGATTAACGGAATCTGCGACAAGCAAACTAACAAGACAATTCTTGATTTCTTGATCGATGACGATAAGATCAACAGAAACTTCATGCAGTTGATCAACGACGATGGCCTGAGCTTCAAGGATATTATCCAAAAGGCTCAGGTTGTGGGAAAGATTGACGATGTTAAGCAAGTTGTGCAGGAGCTTCCAGGCTCTCCTGCCATTAAGAAGGGTATCCTCCAATCAATCAAGATCGTTGATGAACTTGTTAAGGTGATGGGTCATGCTCCAGAGAGCATTGTGATCGAAATGGCAAGGGAGAATCAGACCACTGCTAGAGGAAAGAAGAACTCTCAACAGAGGTACAAGAGAATCGAAGACGCTCTGAAGAACCTCGCCCCAGGCCTCGATTCTAATATCCTTAAGGAGAACCCTACTGACAATATTCAATTGCAGAATGATAGGCTTTTCTTGTACTACCTGCAAAACGGCAAGGACATGTACACAGGAGAGGCCTTGGATATCAACCAACTGTCAAATTACGACATTGATCACATCGTGCCACAGGCATTTATTAAGGACGATTCCCTCGACAATAGGGTTCTTACATCTTCAAAGGACAACAGGGGAAAGTCTGATAATGTGCCTTCAATCGAAGTTGTGCAGAAGAGAAAGGCCTTCTGGCAACAGTTGCTGGATTCTAAGCTCATTTCAGAGAGGAAGTTTAACAATCTCACCAAGGCAGAAAGGGGTGGACTTGACGAGAGAGATAAGGTTGGCTTCATTAAGAGGCAACTGGTGGAGACAAGACAGATCACCAAGCATGTTGCACAAATTCTCGATGCTAGATTTAACACTGAAGTTAACGAGAAGAATCAGAAGATCAGGAAGGTGAAGATTATCACACTCAAGTCCAACCTTGTTAGCAATTTCAGGAAGGAATTTGGCCTTTACAAAGTTAGAGAGATCAATGACTACCATCACGCACACGATGCTTACTTGAACGCCGTTGTGGCTAAGGCCATTCTCAAGAAGTACCCAAAGCTTGAACCTGAGTTCGTGTACGGTGACTACCAAAAGTACGATCTTAAGAGGTACATCTCTAGGTCAAAGGACCCAAAGGAAATTGAGAAGGCAACCGAGAAGTACTTCTTTTACTCAAACCTCCTTAATTTCTTTAAGGAAGAGGTGCATTACGCTGATGGAACTATCATCAAGAGAGAAAACATCGAGTACAGCAAGGACACAGGCGAAATTGCTTGGAACAAGGAGAAGGATTTCGCCACCGTTAGGAAGGTGTTGTCTTGTCCACAAGTTAATATTGTGAAGAAGACTGAAGTTCAGACAGGCGGTTTTTCCAAGGAGAGCATCTTGCCTAAGAGGAACTCCGACAAGCTGATTGCTAGAAAGAAGGACTGGGATCCAAAGAAGTACGGAGGCTTTGATTCCCCTACAGTTGCCTACAGCGTGCTCGTTGTGGCAAAGGTGGAAAAGGGAAAGTCCAAGAAGTTGAAGAGCGTTAAGGAGCTGGTGGGAATTACCATCATGGAAAGATCCAGCTTCGAGAAGGACCCAGTGGATTTTCTTGAAGCCAAGGGATACAAGGAAGTTAGGAAGGATTTGATTATCAAGCTGCCTAAGTACTCCCTTTTCGAATTGGAGAATGGCAGGAAGAGAATGCTCGCAAGCGCTGGTGAACTTCAAAAGGGAAATGAGCTGGCCCTCCCATCTAAGTACGTTAACTTTCTTTACTTGGCATCTCACTACGAGAAGCTTAAGGGTTCACCTGAAGACAACGAGCAAAAGCAGTTGTTCGTGGAACAACATAAGCACTACCTGGATGAAATTATCGAGCAGATCTCTGAGTTTTCAAAGAGGGTGATTTTGGCCGACGCAAATCTCGATAAGGTTCTTTCAGCATACAACAAGCATAGGGATAAGCCAATCAGAGAACAAGCTGAGAATATTATCCACCTGTTCACCCTCACTAACCTTGGTGCTCCTGCAGCTTTCAAGTACTTTGACACAACCATTGATAGGAAGAGATACACATCCACCAAGGAAGTTTTGGACGCCACCCTGATTCATCAGTCTATCACCGGACTTTACGAGACTAGGATCGATTTGTCACAACTGGGTGGAGACAAGAGAACTGCCGATTCCCAGCACAGCACCCCTCCTAAAACTAAACGCAAGGTCGGATCTGGTCCTAAGAAAAAACGTAAAGTGTAG

The c-myc NLS is highlighted in yellow, SV40 NLS sequences are highlighted in green, the *OpenCRISPR-1* sequence codon-optimized for *Triticum aestivum* (primary expression host) and *Medicago sativa* (secondary expression host) is highlighted in blue.
